# AlphaFold2 captures the conformational landscape of the HAMP signaling domain

**DOI:** 10.1101/2023.05.23.541686

**Authors:** Aleksander Winski, Jan Ludwiczak, Malgorzata Orlowska, Rafal Madaj, Kamil Kaminski, Stanislaw Dunin-Horkawicz

**Affiliations:** Laboratory of Structural Bioinformatics, Centre of New Technologies, University of Warsaw, Banacha 2c, 02-097 Warsaw, Poland; Institute of Evolutionary Biology, Faculty of Biology, Biological and Chemical Research Centre, University of Warsaw, Żwirki i Wigury 101, 02-089, Warsaw, Poland; Department of Protein Evolution, Max Planck Institute for Biology Tübingen, Max-Planck-Ring 5, 72076 Tübingen, Germany

## Abstract

In this study, we present a conformational landscape of 5000 AlphaFold2 models of the HAMP domain, a short helical bundle that transduces signals from sensors to effectors in two-component signaling proteins such as sensory histidine kinases and chemoreceptors. The landscape reveals the conformational variability of the HAMP domain, including rotations, shifts, displacements, and tilts of helices, many combinations of which have not been observed in experimental structures. HAMP domains belonging to a single family tend to occupy a defined region of the landscape, even when their sequence similarity is low, suggesting that individual HAMP families have evolved to operate in a specific conformational range. The functional importance of this structural conservation is illustrated by poly-HAMP arrays, in which HAMP domains from families with opposite conformational preferences alternate, consistent with the rotational model of signal transduction. The only poly-HAMP arrays that violate this rule are predicted to be of recent evolutionary origin and structurally unstable. Finally, we identify a family of HAMP domains that are likely to be dynamic due to the presence of a conserved pi-helical bulge. All code associated with this work, including a tool for rapid sequence-based prediction of the rotational state in HAMP domains, is deposited at https://github.com/labstructbioinf/HAMPpred.

## Introduction

The HAMP domain is an approximately 50-amino acid protein domain consisting of two helices and a typically unstructured region connecting them^1–4^. It is mostly found in dimeric signal transduction proteins, such as prokaryotic sensor histidine kinases and chemoreceptors, where it links the sensory extracellular domain to the downstream effector and acts as a signal transduction modulator (Figure 1). HAMP domains are also found in unusual receptors with complex domain compositions, including di– and poly-HAMP arrays in which two or more HAMP domains are concatenated^1,5,6^. In the context of a dimer, the HAMP domain forms a four-helical coiled coil^7^ in which all helices are oriented in parallel (Figure 2). Typical coiled-coil domains are characterized by the presence of the seven-residue (heptad) sequence repeat [abcdefg]_n_, in which residues at positions *a* and *d* face the center of the bundle. These residues from opposite helices interact via the knobs-in-holes packing^7^ to form regular supercoiled structures.

**Figure 1.**
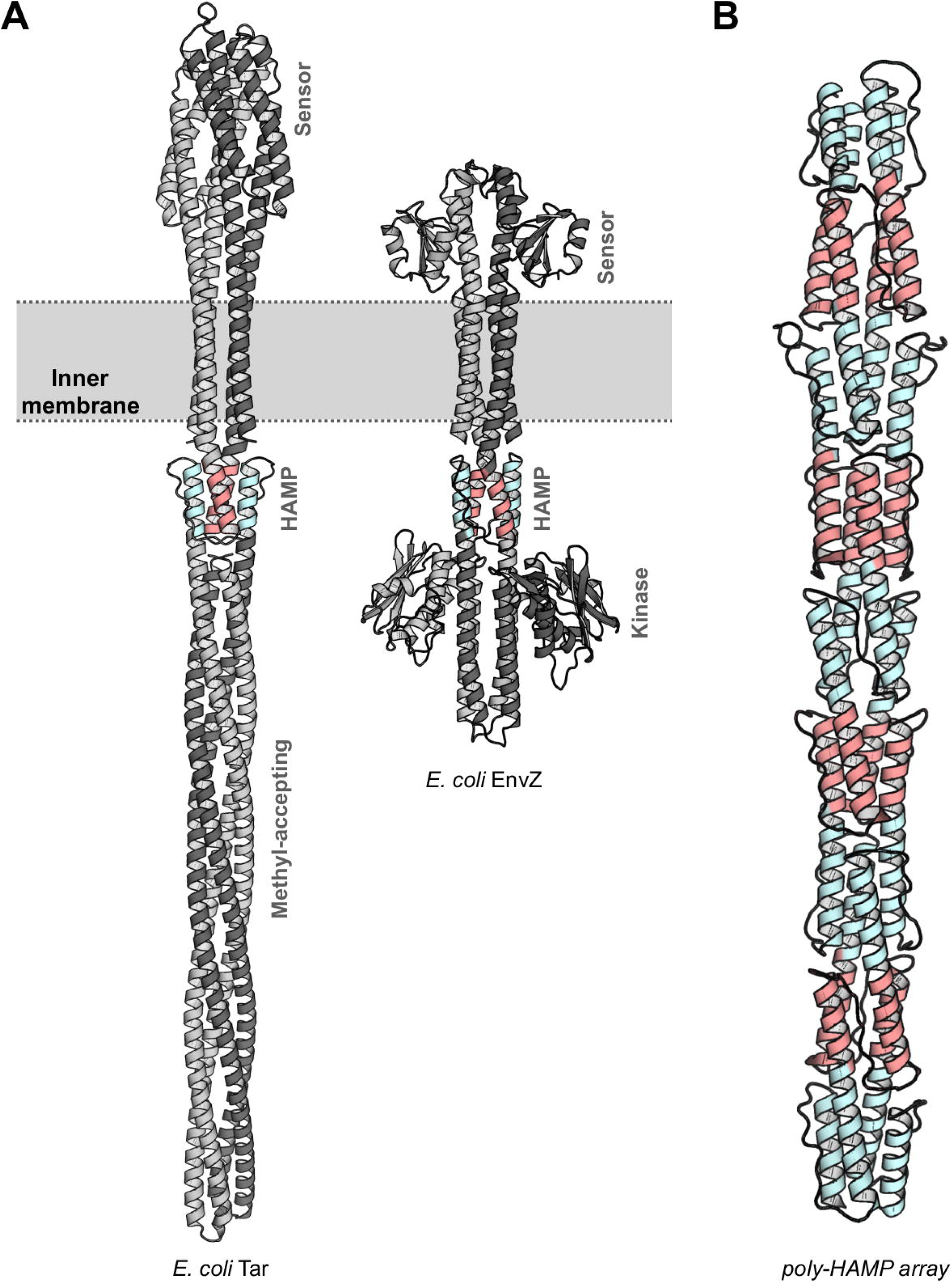
AlphaFold2 models of (A) *Escherichia coli* Tar and EnvZ dimeric receptors (in both models, the HAMP domain is shown in color, with its N– and C-terminal helices shown in red and cyan, respectively) and (B) a poly-HAMP array of a putative receptor (WP_011140637), with its even and odd HAMP domains shown in red and cyan, respectively.

**Figure 2.**
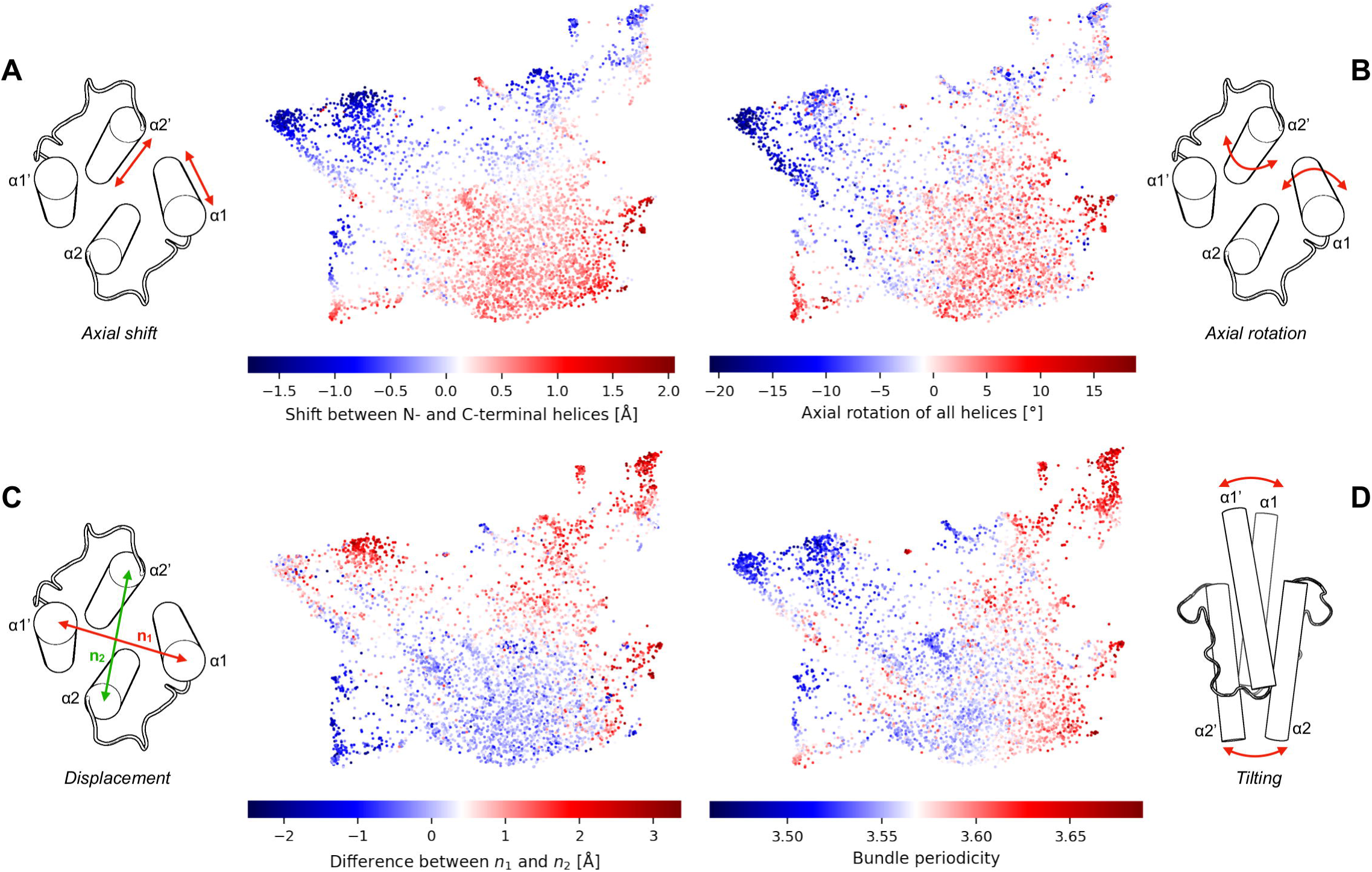
Conformational landscape of 5000 AlphaFold2 HAMP models. Each point corresponds to a single model and their relative positions reflect the global structural similarity measured by FoldSeek. In each panel, the conformational map is colored according to the coiled coil structural parameters measured in each model.

The HAMP domains have been extensively studied both experimentally and computationally to characterize the conformational changes they undergo during signal transduction. These changes, which affect the relative positions of the HAMP helices, can be decomposed into basic motions: axial shift, axial rotation, horizontal displacement, and tilting (Figure 2). For example, the gearbox^8^, piston^9,10^, and scissoring^11^ models describe conformational changes in HAMP domains as coordinated rotations, asymmetric shifts, and tilting of helices, respectively. However, many studies suggest that the changes are better described as combinations of two or more motions^4,12–14^. Another proposed mechanism for HAMP signaling focuses on its dynamics. In this model, the regulation of signaling is attributed to changes in structural stability, rather than a switch between discrete conformations^15–17^. All of these models relating HAMP domain function to its conformation rely on a limited number of experimental structures that may not represent the full conformational landscape accessible to HAMP domains.

AlphaFold2 predictions have been shown to provide insight not only into the static structure of proteins, but also into their conformational plasticity ^18–21^ and dynamics ^22,23^. Considering this, we used it to explore the conformational diversity of the HAMP domain by analyzing models generated for a set of >5000 phylogenetically diverse sequences originating from 81 families defined by sequence similarity^1^. These families have been grouped, also based on sequence similarity, into seven classes (labeled A through G), each containing one or more families, and this nomenclature is used throughout the text. Individually, the obtained models likely represent resting states, which are the most energetically favorable conformations. However, when analyzed as a whole, they provide a comprehensive view of the conformational space accessible to HAMP domains in general.

## Materials and Methods

### AlphaFold2 modelling

6456 aligned HAMP sequences were obtained from our previous work^1^ and clustered at 70% identity with MMseqs2^24^ to remove redundancy, resulting in a set of 5462 sequences. From this set, 74 sequences containing gaps in helices were removed, resulting in a final set of 5388 sequences. The HAMP domains of the final set were modeled as dimers using the ColabFold 1.5.2 environment^25^ with default parameters (no PDB templates were used). In addition, sequences corresponding to 31 experimental structures were modeled for method validation (Table 1 and Supplementary File 1).

### Conformational landscape calculation

All models and 31 experimental structures were compared to each other using FoldSeek^26^ with the following parameters: maximum number of target sequences 6000, E-value cut-off 10, coverage mode 0, alignment type 1 (global alignment), and exhaustive search flag. From the obtained results, the pairwise LDDT (Local Distance Difference Test) scores were extracted and converted into a square distance matrix. The matrix was used as input to the UMAP dimensionality reduction tool^27^ with the following parameters: number of target dimensions 2, number of neighbors 200, and metric “precomputed” (Figure 2).

### Measurement of coiled coil parameters

SamCC Turbo^28^ was used to measure coiled-coil structural parameters in the AlphaFold2 models (Figure 2) and 31 experimental structures. This tool analyzes a given coiled-coil structure by dividing it into layers of *n* Cα atoms (where *n* is the number of helices in a bundle) lying approximately in a plane perpendicular to the bundle axis. The structure parameters are then calculated separately for each layer. In the case of HAMP domains, the N– and C-terminal helices were considered separately, and all measurements were made for 11-residue long segments (see Table 1 for the segments defined for the experimental structures). The segments were defined based on the manually curated multiple sequence alignment^1^, and thus the results are comparable between all HAMP domains analyzed.

The axial shifts of the N– and C-terminal 11-residue helical segments were calculated relative to the bundle axis (Figure 2A). Then, by subtracting the C-terminal shift from the N-terminal shift, a single number was obtained representing the relative shift between the N– and C– terminal helices in a given HAMP domain.

Helical rotation (Figure 2B) was calculated as the difference between the measured Crick angles and those expected for a canonical coiled coil assuming knobs-in-holes packing (Crick angles quantify the orientation of residues with respect to the bundle axis, where 0° indicates a residue pointing directly toward the bundle axis, while the +/-180° indicates a residue pointing directly outward from the bundle axis). The expected values were defined individually for each heptad position as a:19.5°, b:122.35°, c:-134.78°, d:-31.92°, e:70.92°, f:173.78°, and g:-83.35°. Thus, each HAMP model was assigned two 11-element vectors describing the rotation of the N– and C-terminal helices. Values close to zero indicate rotation compatible with knobs-into-holes packing, while values above or below zero indicate da-x or x-da packing, where an additional third residue is co-opted to the hydrophobic core (Figure 3A and C). Global helix axial rotation was defined as the difference between the rotation of the N– and C-terminal helices divided by two (see black solid lines in Figure 3). The 11-element global helix axial rotation vector represents the relative rotation of the N/C-terminal helices. Finally, by averaging this vector, a HAMP rotational state can be represented by a single number ranging from –26° to +26° (Figure 2B).

**Figure 3.**
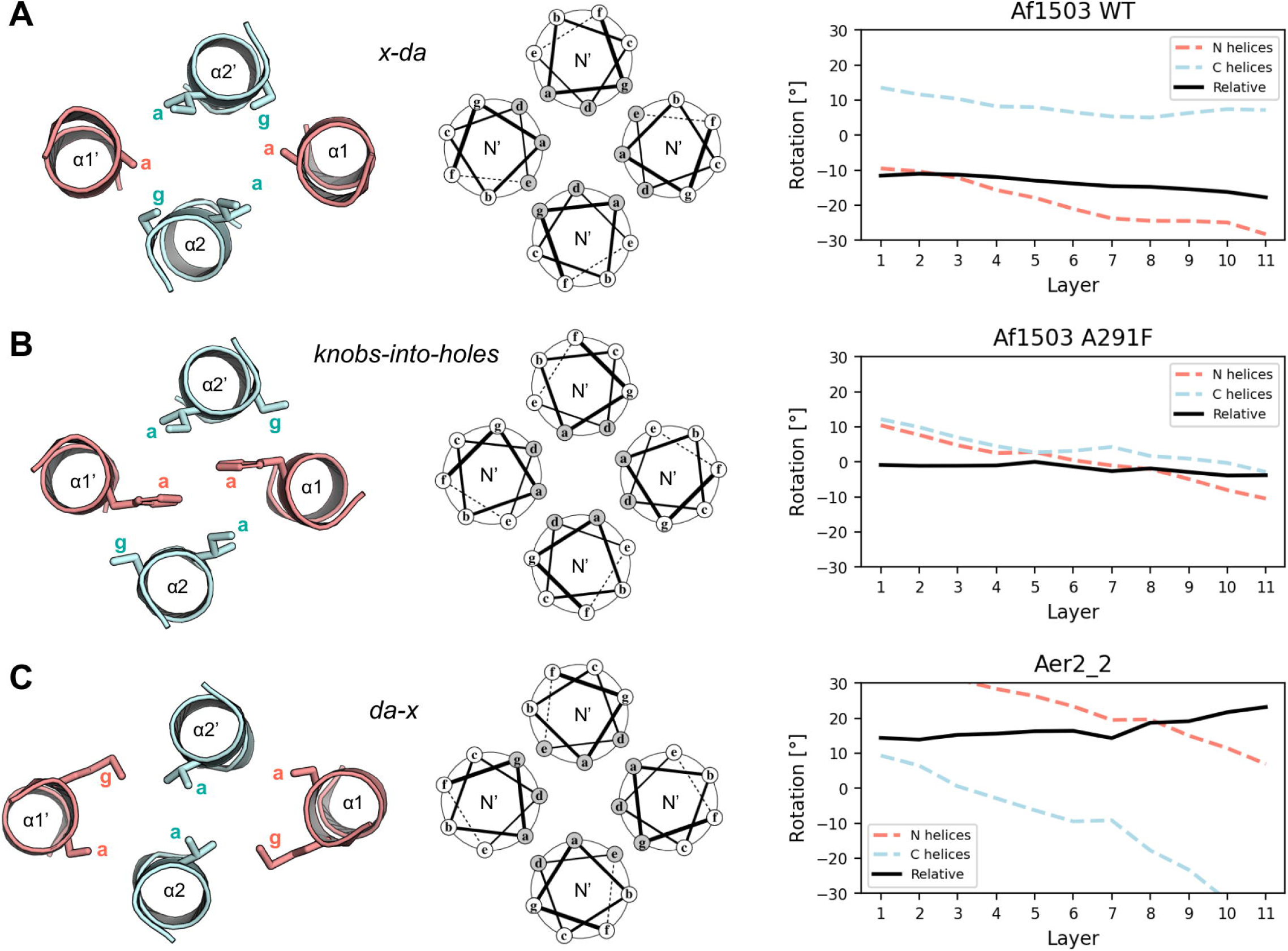
The three basic conformations observed in experimental HAMP structures. (A) The WT Af1503 HAMP domain adopts an x-da packing in which three residues form the hydrophobic core. (B) The A291F variant of the Af1503 HAMP domain adopts the canonical knobs-into-holes packing in which two residues form the hydrophobic core. (C) The second HAMP of the Aer2 protein poly-HAMP array adopts a da-x packing in which three residues form the hydrophobic core. The left panels show cross sections of HAMP domain structures viewed from the N-terminus (linkers between N– and C-terminal helices are not shown), with letters indicating heptad positions. The middle panels show the corresponding wheel diagrams showing which heptad positions are involved in the formation of the hydrophobic core. The right panels show measurements of the axial rotation of the N– (red) and C– terminal (cyan) helices. The black lines indicate the global rotation, calculated as the difference between the rotation of the N– and C-terminal helices (see Methods for details).

To calculate the helix displacement, the average distances between the N– and C-terminal helices in the defined 11-residue segments were first determined (Figure 2C). Then, by subtracting the distance between the C-terminal helices from that of the N-terminal helices, a single value defining the cross-sectional shape was defined.

Finally, the helical tilt (Figure 2D) was defined by calculating the bundle periodicity, which quantifies the degree of supercoiling of the helices. A value of 3.63, corresponding to the periodicity of the undistorted helix, indicates the absence of supercoiling (parallel helices, not tilted), while values lower and higher than 3.63 indicate left– and right-handed supercoiling, respectively (helices tilted).

### HAMPpred prediction model

To enable rapid prediction of the rotational state based on a HAMP sequence, a deep learning model was developed. The set of 5388 HAMP models was filtered by removing models with an absolute mean global rotation greater than 26 or a standard deviation of global rotation greater than 14, resulting in a set of 4972 models. In each model, the sequences of the N– and C-terminal helices (model input) were encoded separately as a matrix of shape (11, 23), where 11 is the helix segment length and 23 is the one-hot encoded amino acid and its hydrophobicity and side-chain size (we used the Kyte and Doolittle hydrophobicity scale^29^ and obtained side-chain sizes from the Jena Library of Biological Macromolecules^30^). The encodings for the N– and C-terminal helices were concatenated along the last dimension, resulting in a matrix of the form (11, 46) for each model. The global helix axial rotation vectors (model output) were encoded using sin and cos functions, resulting in a matrix of the form (11, 2), where 11 is the global helix axial rotation vector length and 2 is the sin and cos of its values. Such encoding avoids periodic effects of degrees and can be inverted using the arc tangent function.

This set of 4972 encoded HAMP sequences (input) and conformations (output) was randomly divided into 3978 training and 994 validation examples. The training set was used to train 4 model variants, while the validation set was used for early termination and selection of the best model architecture. The general model architecture was 3 convolutional filters, each consisting of 3 parallel layers with kernel sizes of 3, 4, and 7 and a batch normalization layer, 1 or 2 bidirectional LSTM (Long Short-Term Memory) layers, 1 or 3 dense layers, and an output layer. Each model was trained for 100 epochs with an initial learning rate of 0.0001. Tanh was used as the activation function and mean square error (MSE) as the loss function. If the validation loss did not improve for 7 epochs, the learning rate was set to a value 10 times lower than the previous one. If the validation loss did not improve for 15 epochs, training was stopped. The best model had two LSTM layers and a single dense layer.

## Results and discussion

We modeled over 5000 HAMP sequences using AlphaFold2, which represent 81 families identified in our previous work^1^. These sequences come from different domain contexts, ranging from those found in canonical sensor kinases and chemoreceptors to unusual arrangements involving the formation of di– and poly-HAMP arrays. All obtained models were compared with each other, and the resulting similarity matrix was visualized as a 2D conformational landscape (see the Methods for details). To highlight structural differences, we color-coded the conformational map based on the coiled-coil parameters measured in each HAMP model. Specifically, we quantified the relative orientation of the helices in terms of axial shift, rotation, displacement, and tilting (Figure 2).

The colored maps show overlapping, partially correlated (ρ=0.55) gradients of helix axial shift (−1.7 Å to 2 Å) and rotation (−21° to 19°; Figure 2A and B). Such coordinated screw-like motion combining axial rotation and shift, is compatible with the rotational model of signal transduction by HAMP domains^8^ and was also observed in the sensory domain of Tar protein, a chemoreceptor containing the HAMP domain^31^. Analysis of coiled coils similar to HAMP domains suggests that the observed shift is due to the constraints imposed on the side chains by a given rotational state^7,32^. Effectively, adopting extreme rotation values requires adjusting the relative shift between the helices. However, this shift does not disrupt the coiled coil register and is symmetrical, making it incompatible with the piston model^9,10,33^, which assumes an asymmetric, register-disrupting motion that can propagate to adjacent domains.

The remaining two parameters, displacement (Figure 2C) and tilt (Figure 2D), show no correlation to each other or to rotation and shift. The horizontal helix displacement, defined as the difference in distance between the N– and C-terminal helices, ranges from –2.5 Å to 3.4 Å, with values of 0 Å indicating a perfectly square cross-section. Motions based on such displacements have been suggested by molecular dynamics simulations^34^. The tilt, described by the bundle periodicity coiled-coil parameter, ranges from 3.45 (left-handed bundle with strong helical tilt) to 3.66 (slightly right-handed bundle with nearly parallel helices). Such differences in helical tilt have been described in HAMP domains and shown to be associated with rotation^4,12^.

The experimental HAMP structures are distributed along the helical rotation and shift gradients, with the HAMP domains of Af1503 and the second HAMP of the Aer2 poly-HAMP array defining the boundaries (Figure 4). Notably, the structural variability among the HAMP variants is very limited. For example, although Af1503 mutants differ in both structure and activity of downstream catalitic domain^35^, they are localized in a small area of the conformational landscape. Similarly, other HAMPs for which mutated and WT structures are available, i.e. NarQ and Aer2 (HAMP1) are also grouped together. This suggests that HAMP domains operate within specific conformational ranges, albeit sufficient to transduce the signal, and that these ranges may differ substantially among HAMP domains. It is also worth noting that most of the experimental HAMP structures were determined in the context of other domains and/or mutations. The fact that they cluster together with the AlphaFold2 models suggests that the landscape captures such induced conformations. Finally, the results obtained show that while the available experimental structures sample the full range of axial rotation and shift states, there are large areas of the conformational landscape that are not covered.

**Figure 4.**
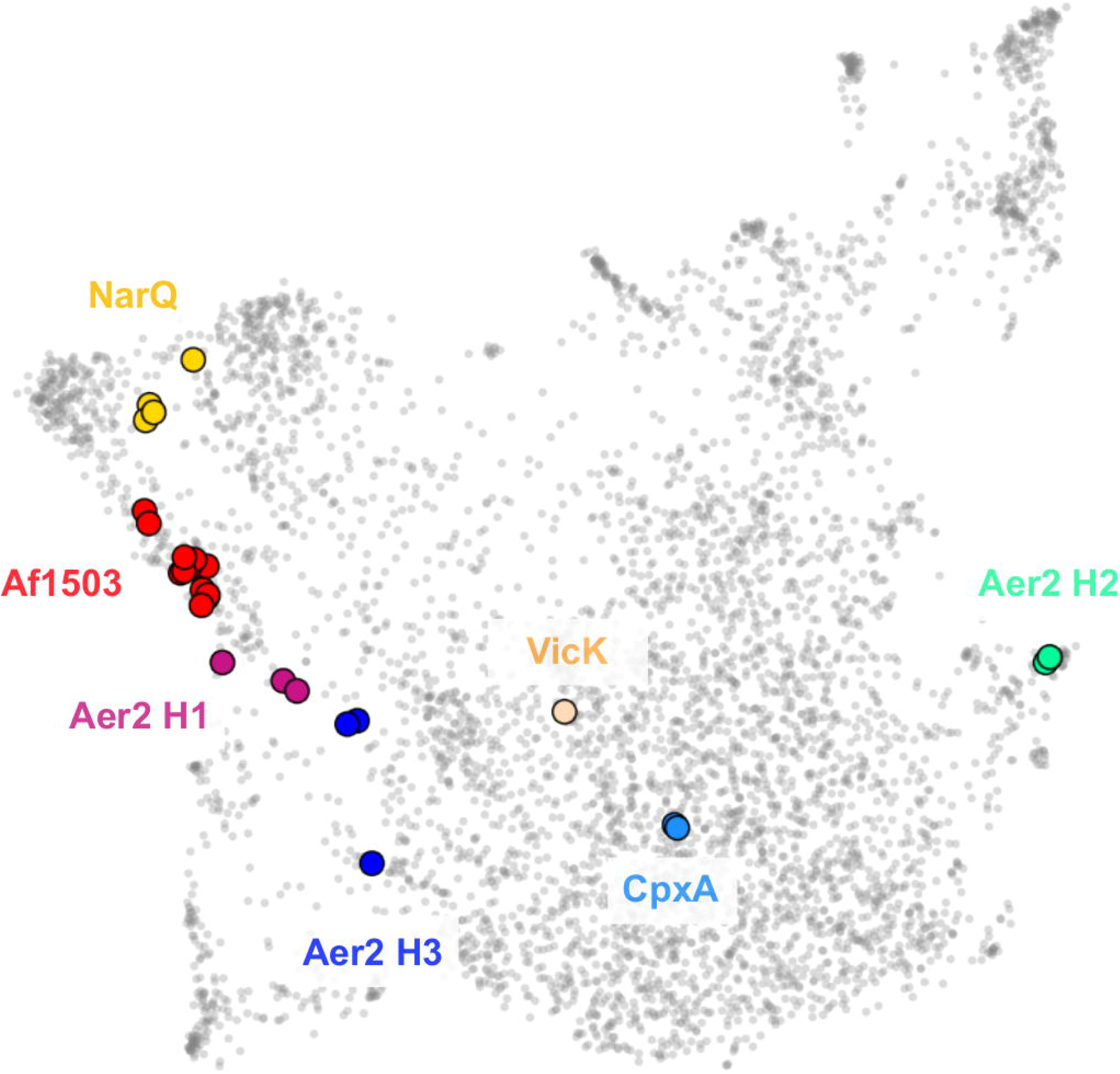
Localization of 31 experimental HAMP structures in the calculated conformational landscape. Gray dots indicate AlphaFold2 models, while experimental structures are shown in color. Each color corresponds to a single HAMP domain type and its variants.

We observed the same pattern of structural conservation in HAMP families defined by sequence similarity^1^. In many of them, the HAMP domains show a preference for a resting conformation, even when they share a low sequence identity (Figure 5A, values in brackets; see also Supplementary File 2). We hypothesize that such a conserved preference for the resting conformation reflects the “tuning” of a given HAMP domain (or HAMP family) to function in the particular context of other domains, and that these domains co-evolve to maintain a balance between their conformational preferences.

**Figure 5.**
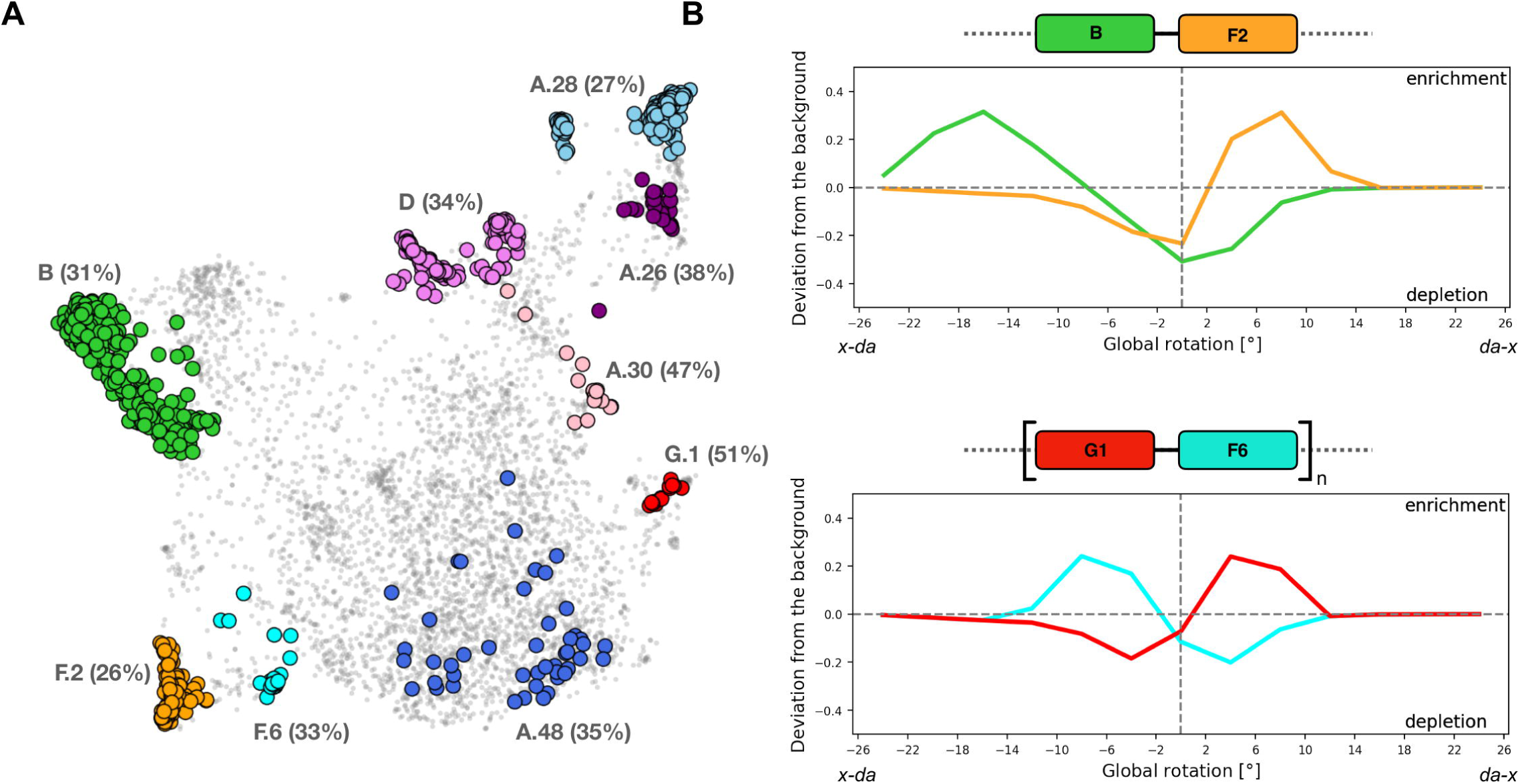
Localization of HAMP families in the calculated conformational landscape. (A) Selected HAMP families, defined on the basis of sequence similarity, are shown in color. Numbers in brackets indicate the average sequence identity within a family. (B) Conformational preferences in di– and poly-HAMP arrays. A deviation from the baseline (0 on the y-axis) indicates that a particular HAMP family is depleted or enriched in certain conformational states. Family B HAMP domains occur alone or in combination with family F2 HAMP domains to form di-HAMP modules. Family G1 and F6 HAMPs exclusively form poly-HAMP arrays in which the G1-F6 module can be repeated several (n) times.

To investigate this possibility, we analyzed unusual HAMP families that occur exclusively together and from di– and poly-HAMP arrays in which the two different families alternate^1,5^. This alternating pattern can be explained by the rotational model, which describes the structural changes that occur in a HAMP domain as a coordinated rotation of its helices between two boundaries^8^. For the well-studied HAMP domain of the *Archaeoglobus fulgidus* Af1503 protein, the rotation boundaries are defined by two states: the canonical knobs-into-holes packing, in which two residues form the hydrophobic core (Figure 3B)^35^, and an alternative form of packing, the x-da packing, in which an additional residue is co-opted into the core (Figure 3A)^8^. The transition from the canonical to the x-da packing is achieved by rotating the N– and C-terminal helices counterclockwise and clockwise, respectively, relative to the knobs-in-holes packing^35^ (compare Figure 3A and B). If a HAMP domain similar to the Af1503 HAMP were to function in the context of another HAMP domain, the latter would have to be capable of receiving reversed rotational signal from the former one. This can be achieved by adopting two conformations: knobs-into-holes (as in the case of Af1503) and an alternative one, da-x (rotation of +26°), observed for example in the second HAMP of Aer2 (Figure 3C), in which the helices are rotated in the opposite direction (N– and C-terminal helices are rotated clockwise and counterclockwise, respectively, relative to the knobs-in– holes packing).

To assess the conformational preferences of HAMP domains from di– and poly-HAMP arrays, a normalized distribution (52 bins ranging from –26° to +26°) of rotation values across all 5000 models was first computed. This baseline distribution was then subtracted from the analogous distributions calculated for the families of interest (Figure 5B), and the differences were plotted, indicating rotational states that are more (enrichment) or less (depletion) common. The results obtained suggest that the conformational preferences of adjacent HAMP domains of di-HAMP and poly-HAMP arrays are opposite. For example, some of the B family HAMPs are followed by a HAMP from another family, F2, to form a di– HAMP module. The B HAMPs show a preference for x-da packing; however, the F2 HAMPs show the opposite preference, favoring da-x packing. In such B-F2 modules, the B-HAMPs would rotate between x-da and knobs-into-holes conformations, whereas the F2-HAMPs would rotate between da-x and knobs-into-holes conformations (Figure 3). The localization of these families in the conformational landscape (Figure 2) shows that rotation is also accompanied by displacement and tilting of the helices, suggesting a more complex nature of the conformational transitions.

While B-type HAMPs, mentioned above, are found both alone and in association with F2– type HAMPs, the G1 and F6 HAMP domains almost exclusively form long poly-HAMP arrays in which sequences from both groups alternate (Figure 1B). The results obtained (Figure 5B) indicate that HAMP domains from these two groups also have clearly opposite preferences for the resting state conformation, where the F6 family HAMPs are shifted to x-da packing and the G1 family HAMPs are shifted to da-x packing. This alternating pattern is also observable in the experimental structure of 3-HAMP array of the Aer2 receptor^4^, where HAMP1 and HAMP3 occupy quite similar positions on the map (x-da packing; Figure 4), whereas HAMP2 localized between them can be found on the opposite site of the map (da-x packing).

We propose that the conserved conformational preferences of the HAMP families could be used to aid in the design of hybrid proteins in which a HAMP domain is placed in a new context (e.g., upstream of another output domain or in conjunction with other HAMPs). We also envision that analogous compatibility rules could be derived for other coiled-coil domains of two-component signaling systems.

Finally, we investigated how the observed HAMP conformations might affect structural stability, a property that has been proposed to influence the signaling process^15–17^. Previous research has shown a correlation between AlphaFold2 scores and protein dynamics as assessed by NMR^22^ and molecular dynamics^23^. Building on this insight, we calculated the average pLDDT and PAE for each of the 81 HAMP families (Figure 6) and found that only a few of them exhibited persistently decreased scores. Interestingly, most of these potentially unstable HAMP families are part of unique poly-HAMP arrays, where HAMP domains from the same family are repeated. These arrays are not only rare, but also different from the patterns seen in other poly-HAMPs, where HAMP families with opposite resting conformations alternate. We hypothesize that this predicted instability may represent an evolutionary intermediate in the formation of poly-HAMP arrays. In this scenario, these arrays may not result from the amplification of the entire di-HAMP module (as in cases such as G1-F6), but rather from a single domain. Consequently, in the view or rotational model, the resulting array may remain unresponsive unless the constituent HAMP domains have greater freedom in terms of accessible conformations, a feature that increased dynamics can provide. The notion that this type of single-family amplification may represent an evolutionary intermediate is further supported by the remarkable similarity between HAMP domains within these atypical poly-HAMP arrays. For example, in the five array-forming F7 HAMPs of a putative receptor (ZP_01998985), the average similarity reaches 70%, suggesting that it likely originated from a relatively recent evolutionary event.

**Figure 6.**
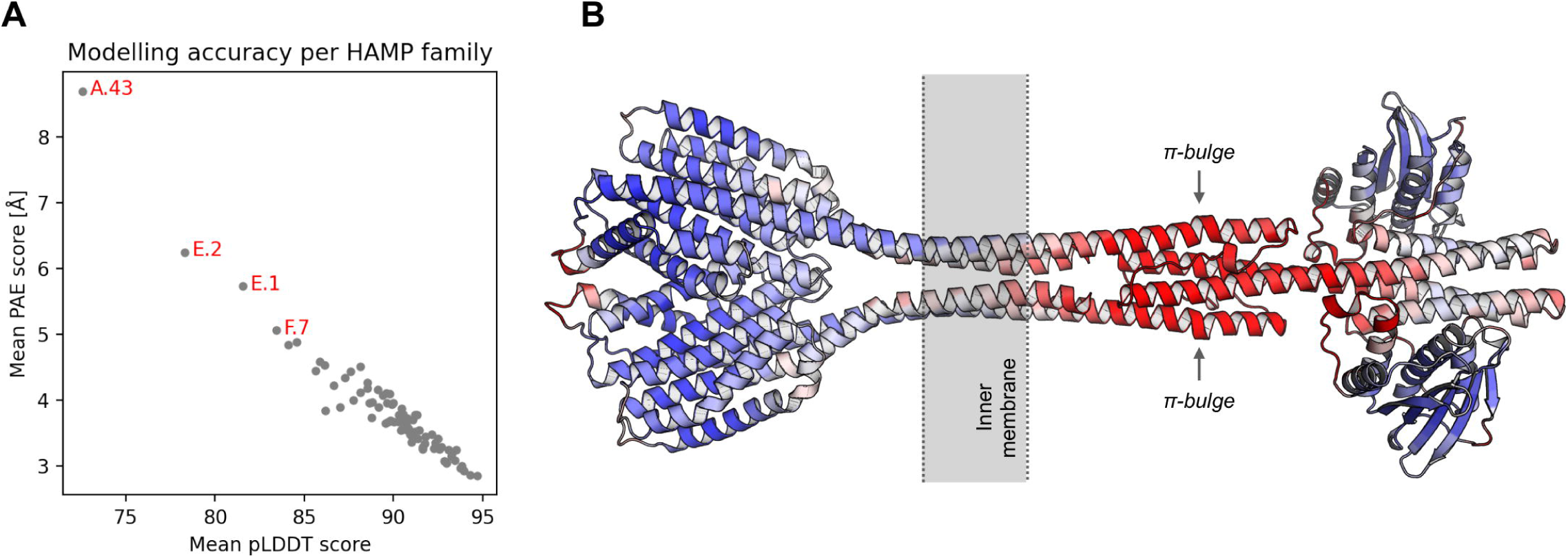
Modeling accuracy across 81 HAMP families. (A) Each point represents the average pLDDT and PAE AlphaFold2 scores of models from a single HAMP family. Four families with the worst scores, indicating possible structural instability, are highlighted. (B) AlphaFold2 model of the entire receptor (WP_011030628) containing HAMP from family A43. The structure is colored according to the pLDDT score, with red indicating the worst score and blue the best.

Among the HAMP families predicted to be the least stable, A43 is the only exception as it doesn’t participate in poly-HAMP formation. These HAMPs are found in a single copy in the context of an extracellular sensing domain and an output kinase domain. By modeling the structure of the entire receptor containing an A43 HAMP (WP_011030628), we discovered some peculiarities (Figure 6B). In the model, only the HAMP domain shows low pLDDT scores, indicating its structural instability. This instability could be attributed to the presence of pi-bulge, a secondary structure element known for its destabilizing effects^36,37^, and also studied in the context of coiled coils^38^. The pi-bulge observed in the model are also detectable at the sequence level by PiPred^39^, suggesting that this may be a conserved feature of the A43 family. In addition, A43 HAMPs have unusually long linkers between the helices that don’t twist around but instead allow for the formation of a much longer HAMP domain. This family may serve as a model to study the role of dynamics in signal transduction through the HAMP domain.

As described above, the AlphaFold2 models provide insight into the conformational landscape of HAMP domains; however, their applicability can be limited by the time required to run folding simulations (approximately 1-2 minutes per HAMP structure). To address this problem, we used AlphaFold2 structures to train a prediction model, HAMPpred, which is capable of predicting the resting rotational state for thousands of HAMP sequences within seconds. The benchmark with 31 experimental structures (Table 1) showed that HAMPpred provides sufficient accuracy to discriminate between the extreme rotational states (Figure 7A). AlphaFold2 performs better in modeling fine details of the HAMP domains (Figure 7B), with only two examples where the models and experimental structures differ significantly, i.e., Af1503 HAMP, where the alanine at position 291 was replaced by phenylalanine, and Aer2 HAMP1, where the leucine was replaced by histidine. The difficulty in modeling these cases is probably related to the fact that the phenylalanine and histidine at these positions are never observed in the Af1503 HAMP and Aer2 HAMP1 homologs, respectively.

**Figure 7.**
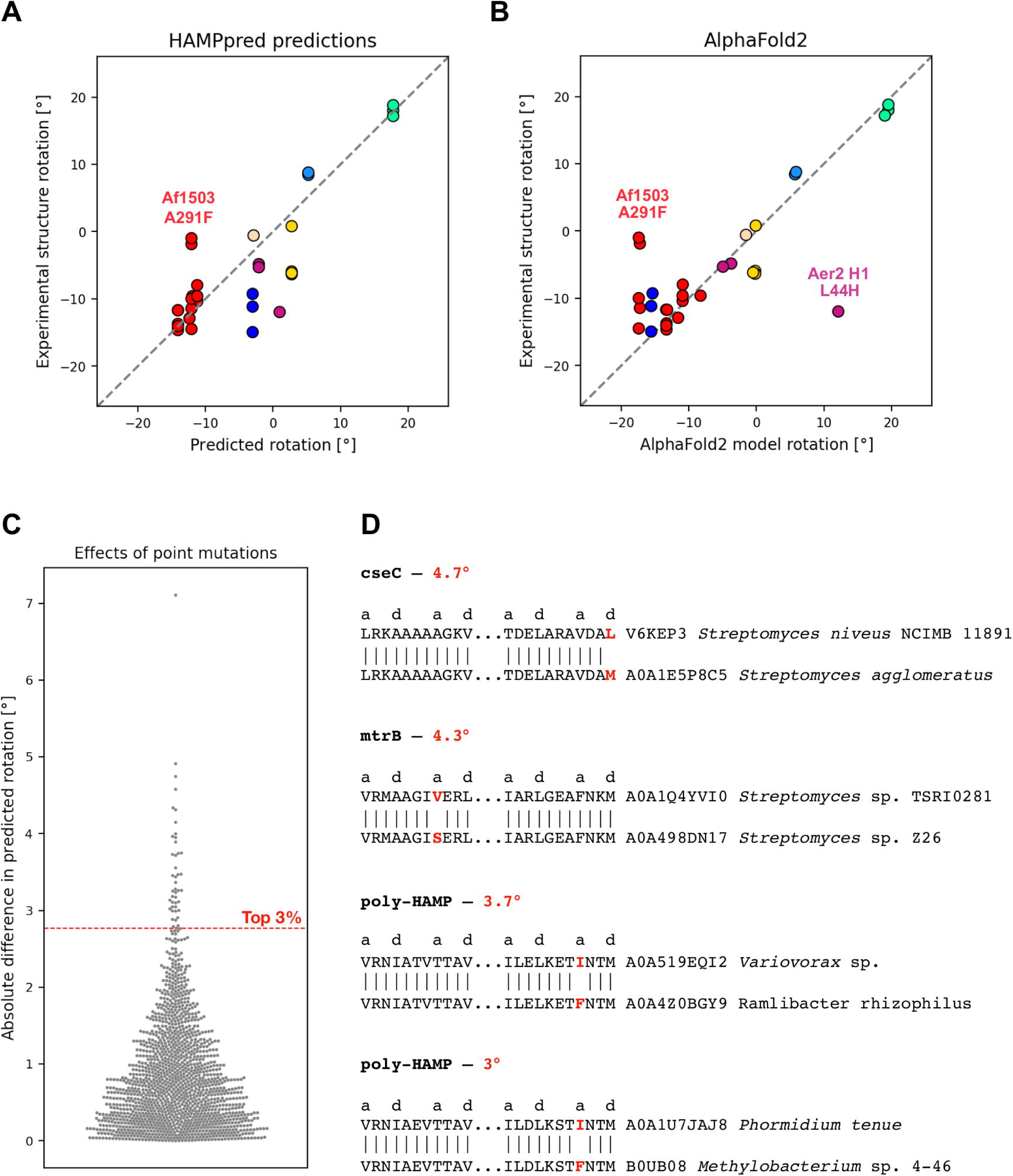
Prediction of the rotational state of HAMP domains. (A) Sequence-based predictions. The x and y axes show the averaged global rotation predicted by HAMPpred and the averaged global rotation measured in the experimental structures, respectively. (B) AlphaFold2 predictions. The x and y axes show the averaged global rotation in the AlphaFold2 models and the experimental structures, respectively. (C) Predicted effects of point mutations in the hydrophobic core of the HAMP domains. Each point represents a pair of HAMP domains that differ by only one residue. The position of each point along the y-axis indicates the absolute difference between their predicted rotational states. The red horizontal line represents the top 3% of these differences. (D) Examples of the top 3% of HAMP domain pairs. Mutations are shown in red.

There is considerable experimental evidence that the introduction of point mutations in HAMP domains can significantly affect the functionality of the entire receptor in which they occur ^8,12,35,40–42^. We hypothesized that such changes may also occur naturally as a result of adaptive evolution, where a single residue change adjusts the preferred conformation of the HAMP domain. To address this question, we used HAMPpred to analyze 1500 pairs of HAMP sequences from the Pfam family PF00672 that differ by only one amino acid at one of the hydrophobic core positions. For all such pairs, we calculated the absolute difference in their predicted rotational states (Figure 7C). For most (>90%) pairs, the predicted difference was less than 2°, consistent with the observation that the resting conformation is conserved in HAMP families. However, in some cases the difference was larger, up to 7°. We examined those with differences in the top 3% and identified a pair of sensor kinases from *Streptomyces niveus* and *Streptomyces agglomeratus* that are putative homologs of CseC, a sensor kinase involved in the response to cell envelope damage in *Streptomyces coelicolor*^43^ (Figure 7D). Another example also comes from the genus Streptomyces, where we identified a pair of kinases from *Streptomyces* sp. TSRI0281 and *Streptomyces* sp. Z26 that are homologs of MtrB, a receptor kinase involved in the regulation of antibiotic production in *Streptomyces coelicolor* and *Streptomyces* venezuelae ^44,45^ (Figure 7D).

The pronounced effects of point mutations were also observed in HAMP domains, such as those of family F6 (Figure 5), that are part of the poly-HAMP arrays (the divergent HAMP domains, such as those of family G1, were not analyzed because they are not covered by the Pfam PF00672 family). In such poly-HAMP arrays (Figure 1B), the initial and terminal HAMP domains are typically distinct from each other and from those in between. In contrast, the middle HAMP domains are very similar to each other, suggesting that they have recently evolved by amplification. For example, the middle HAMP domains of the poly-HAMP array of a putative receptor from *Methylobacterium* sp. (B0UB08) share more than 90% sequence identity on average. The presence of such putative conformation-changing mutations exclusively in the middle HAMP domains may be related to the recovery of conformational equilibrium after amplification of di-HAMP modules in the middle of the array.

In summary, we have shown that AlphaFold2 is a powerful tool for analyzing protein structures that adopt different conformations. AlphaFold2 not only reproduces the conformations observed in experimental structures, but also allows the conformational landscape to be mapped for the entire protein family. The AlphaFold2-based procedure was applied to the HAMP domain family and yielded results consistent with signaling models based on helical rotation, displacement, shift, and tilt, but suggested that the extent of these motions may be specific to individual HAMP families.

## Acknowledgments

This work was supported by „Diamentowy Grant” DI2018010748 (to A.W.) from the Polish Ministry of Science and Higher Education. S.D-H. was supported by institutional funds from the Max Planck Society.

## Authors’ Contribution

Aleksander Winski: conceptualization; investigation; methodology; software; writing – original draft, funding acquisition. Jan Ludwiczak: conceptualization; investigation; methodology. Malgorzata Orlowska: investigation; writing – original draft. Rafal Madaj: investigation. Kamil Kaminski: investigation. Stanislaw Dunin-Horkawicz: conceptualization; methodology; software; writing – original draft; supervision.

