## Supplementary material for "AlphaFold2 captures the conformational landscape of the HAMP signaling domain": Table 1

**Tables**

**Table 1.** Experimental HAMP structures showing different types of hydrophobic core packing. Numbers indicate start and end of N- and C-terminal helices in PDB numbering. Letters above the sequences describe the heptad positions for the canonical knobs-into-holes packing (see Figure 3B).

| **PBD ID** | **N start** | **N sequence** | **N end** | **C start** | **C sequence** | **C end** |
| --- | --- | --- | --- | --- | --- | --- |
|  |  | *a d a d* |  |  | *a d a d* |  |
| 2l7h | 284 | IIELSNTADKI | 294 | 312 | IGILAKSIERL | 322 |
| 2l7i | 284 | IIELSNTFDKI | 294 | 312 | IGILAKSIERL | 322 |
| 2lfr | 284 | IIELSNTADKI | 294 | 312 | IGILAKSIERL | 322 |
| 2lfs | 284 | IIELSNTFDKI | 294 | 312 | IGILAKSIERL | 322 |
| 2y0q | 284 | IIELSNTCDKI | 294 | 312 | IGILAKSIERL | 322 |
| 2y20 | 284 | IIELSNTIDKI | 294 | 312 | IGILAKSIERL | 322 |
| 2y21 | 284 | IIELSNTVDKI | 294 | 312 | IGILAKSIERL | 322 |
| 3lnr | 115 | KMKVVSVVTAY | 125 | 141 | KAQITEAIDGV | 151 |
| 3lnr | 14 | ADRIATLLQSF | 24 | 41 | YERLYDSLRAL | 51 |
| 3lnr | 69 | EAGLAEMSRQH | 79 | 97 | AARIAKGVNEL | 107 |
| 3zcc | 284 | IIELSNTADKI | 294 | 312 | IGILAKSIERL | 322 |
| 3zrv | 284 | IIELSNTFDKI | 294 | 312 | IGILAKSIERL | 322 |
| 3zrw | 284 | IIELSNTVDKI | 294 | 312 | IGILAKSIERL | 322 |
| 3zrx | 284 | IIELSNTFDKI | 294 | 312 | IGILAKSIERL | 322 |
| 3zx6 | 8 | IIELSNTVDKI | 18 | 36 | IGILAKSIERL | 46 |
| 4biu | 190 | ARKLKNAADEV | 200 | 218 | FLAAGASFNQM | 228 |
| 4biv | 190 | ARKLKNAADEV | 200 | 218 | FLAAGASFNQM | 228 |
| 4cq4 | 284 | IIELSNTADKI | 294 | 312 | IGILAKSIERL | 322 |
| 4cti | 284 | IIELSNTFDKI | 294 | 312 | IGILAKSIERL | 322 |
| 4gn0 | 254 | IIELSNTADKI | 264 | 282 | IGILAKSIERL | 292 |
| 4i3m | 115 | KMKVVSVVTAY | 125 | 141 | KAQITEAIDGV | 151 |
| 4i3m | 14 | ADRIATLLQSF | 24 | 41 | YERHYDSLRAL | 51 |
| 4i3m | 69 | EAGLAEMSRQH | 79 | 97 | AARIAKGVNEL | 107 |
| 4i44 | 115 | KMKVVSVVTAY | 125 | 141 | KAQITEAIDGV | 151 |
| 4i44 | 14 | ADRIATLLQSF | 24 | 41 | YERLYDSLRAL | 51 |
| 4i44 | 69 | EAGLAEMSRQH | 79 | 97 | AARIAKGVNEL | 107 |
| 4i5s | 41 | VKQLNAKVRSL | 51 | 68 | LSELVNNVNDL | 78 |
| 5iji | 180 | LNQLVTASQRI | 190 | 208 | LGLLAKTFNQM | 218 |
| 5jef | 180 | LNQLVTASQRI | 190 | 208 | LGLLAKTFNQM | 218 |
| 5jgp | 180 | LNQLVTASQRI | 190 | 208 | LGLLAKTFNQM | 218 |
| 6yue | 180 | LNQLVTASQRI | 190 | 208 | LGLLAKTFNQM | 218 |
